# Evaluation of the 7 & 7 Synch and 7-day CO-Synch + CIDR ^®^ protocols for estrus synchronization of beef cows prior to fixed-time artificial insemination with conventional or sex-sorted semen

**DOI:** 10.1101/2020.08.25.266783

**Authors:** C.M. Andersen, R.C. Bonacker, E.G. Smith, C.M. Spinka, S.E. Poock, J.M. Thomas

## Abstract

An experiment was designed to compare the recently developed 7 & 7 Synch and the 7-day CO-Synch + CIDR protocols for synchronization of estrus among beef cows prior to fixed-time artificial insemination (FTAI) with conventional or sex-sorted semen. *Bos taurus* cows (n = 1538) were blocked based on age and days postpartum (DPP) and randomly assigned to protocol and semen type. Cows treated with the 7-day CO-Synch + CIDR protocol (n = 769) received administration of gonadotropin-releasing hormone (GnRH; 100 μg gonadorelin) and insertion of a 1.38 g intravaginal progesterone releasing insert (CIDR) on Day -10, and administration of prostaglandin F_2α_ (PG; 500 μg cloprostenol) coincident with CIDR removal on Day -3. Cows treated with 7 & 7 Synch (n = 769) received PG and insertion of CIDR on Day -17, GnRH on Day -10, and PG coincident with CIDR removal on Day -3. Estrus detection aids (Estrotect™) were applied to all cows on Day -3, and activation status was recorded at FTAI. Cows received FTAI 66 h after CIDR removal with either conventional (20 × 10^6^ cells per unit) or sex-sorted (4 × 10^6^ cells per unit; SexedULTRA 4M™) semen. A greater proportion of cows expressed estrus (*P* = 0.01) prior to FTAI following 7 & 7 Synch (82%; 629/769) compared with the 7-day CO-Synch + CIDR (64%; 492/769). Estrus expression was also affected by protocol × DPP (*P* = 0.0004), with 7 & 7 Synch resulting in a greater increase in the proportion of cows expressing estrus prior to FTAI among cows with greater DPP. Across protocols, body condition score (BCS) affected (*P* = 0.05) estrus expression, with greater proportion of cows expressing estrus prior to FTAI at greater BCS. Sex-sorted semen resulted in reduced (*P* < 0.0001) pregnancy rates to FTAI. Irrespective of semen type, greater (*P* = 0.001) pregnancy rates to FTAI were obtained among cows treated with 7 & 7 Synch (conventional semen: 72% [280/389]; sex-sorted semen: 52% [199/380]) compared with the 7-day CO-Synch + CIDR (conventional semen: 60% [231/383]; sex-sorted semen: 44% [171/386]). In summary, 7 & 7 Synch resulted in an increased proportion of cows expressing estrus prior to FTAI and an increased pregnancy rate to FTAI with conventional and sexed semen. With these results and ease of application, 7 & 7 Synch offers potential as a platform to improve success with fixed-time AI in beef cows.

## 1. Introduction

Fixed-timed artificial insemination (FTAI) and estrus synchronization are widely applicable reproductive technologies that allow producers to improve economic returns and production efficiency in their operations [1]. Success of these technologies has improved substantially over the last two decades, yielding many benefits with respect to reproductive performance. The increased efficiency and convenience of estrus synchronization and FTAI has facilitated the use of other technologies by beef producers, such as sex-sorted semen and genomic-enhanced EPDs for sire selection [2,3]. In spite of these opportunities, ineffective control of the estrous cycle early in the protocol can present other challenges, such as induced ovulation of physiologically immature follicle at the time of FTAI and reduced fertility [4].

Many short-term estrus synchronization protocols developed for beef cows, such as the commonly used 7-day CO-Synch + CIDR, utilize exogenous gonadotropin-releasing hormone (GnRH) at the start of the protocol in order to induce ovulation of a dominant follicle and initiate a new follicular wave. However, stage of follicular development is a major determinant for an animal’s ability to respond to exogenous hormone treatment during an estrus synchronization protocol [5], and GnRH is only effective for inducing ovulation of a dominant follicle of sufficient physiological maturity. Administration of GnRH at a random stage of the estrous cycle has been observed to induce ovulation with variable success rates, resulting in ovulation in approximately 65% of cows on average [6] and considerably poorer rates possible in any one group. Cows that fail to ovulate are in a stage of follicular development that does not have the capacity to respond to GnRH. The resulting lack of uniformity among cows in subsequent stage of cycle constitutes a potential limitation to pregnancy rates achieved among cows when performing FTAI, as variation in follicular maturity, estrus expression, and timing of ovulation at the end of the protocol can contribute to suboptimal fertility.

Presynchronization treatments to manage stage of cycle in advance of GnRH are one method to address the problem of variation at the start of a protocol, with presynchronization now a widely accepted practice in the dairy industry [7]. However, the beef industry has not readily adopted methods of presynchronization due to the greater number of handling events required. To address the challenge of variable ovulatory response to GnRH, the 7 & 7 Synch protocol incorporates a simple, one-step approach to increase the proportion of cows presenting with a physiologically mature, LH-responsive follicle at the time of GnRH administration [8,9]. By administering prostaglandin F_2 a_ and placing a progesterone-releasing intravaginal insert (CIDR) in advance of GnRH administration, an increased proportion of cows present with a physiologically mature follicle at the time of GnRH administration [8,10,11]. In comparison with cows receiving the 7-day CO-Synch + CIDR protocol, cows receiving 7 & 7 Synch presented with significantly greater largest follicle diameter at the time of GnRH administration, with subsequent CL status and estrus expression suggesting a high ovulatory response to GnRH [8]. In addition, a subsequent large multi-location embryo transfer field trial observed a greater proportion of recipient cows expressed estrus and became pregnant to embryo transfer when treated with 7 & 7 Synch [9]. Based on these observations, the following experiment was designed to evaluate field fertility when using the 7 & 7 Synch protocol prior to FTAI. We hypothesized that the 7 & 7 Synch protocol would result in an increased proportion of cows expressing estrus prior to FTAI, as well as increased pregnancy rates to FTAI with conventional and/or sex-sorted semen when compared with the 7-day CO-Synch + CIDR protocol.

## 2. Materials and Methods

All experimental procedures were approved by the University of Missouri Animal Care and Use Committee.

### 2.1. Animals and Estrus Synchronization

The 7-day CO-Synch + CIDR and 7 & 7 Synch protocols (Figure 1) were used for estrus synchronization among suckled beef cows (n = 1538) of varying age and parity across multiple locations (n = 11) in Missouri and South Dakota in the spring and fall breeding seasons of 2019. Within each location, cows were blocked based on age, days postpartum (DPP), and body condition score (BCS; 1 to 9 scale; 1 = emaciated and 9 = obese) [12] and randomly assigned to treatment within block. Cows treated with the 7-day CO-Synch + CIDR protocol (n = 769) received administration of 100 μg gonadorelin acetate (GnRH; Fertagyl, Merck Animal Health, Madison, NJ) and insertion of a 1.38 g intravaginal progesterone releasing insert (CIDR^®^ Zoetis, Madison, NJ) on Day -10, and administration of 500 μg cloprostenol sodium (PG; Estrumate, Merck Animal Health, Madison, NJ) coincident with removal of CIDR on Day -3. Cows treated with the 7 & 7 Synch protocol (n = 769) received administration of PG and insertion of CIDR on Day -17, administration of GnRH on Day -10, and administration of PG coincident with removal of CIDR on Day -3. For both protocols, all cows received administration of GnRH at the time of FTAI, which was performed on Day 0 at 66 h after CIDR removal and PG administration.

**Figure 1.**
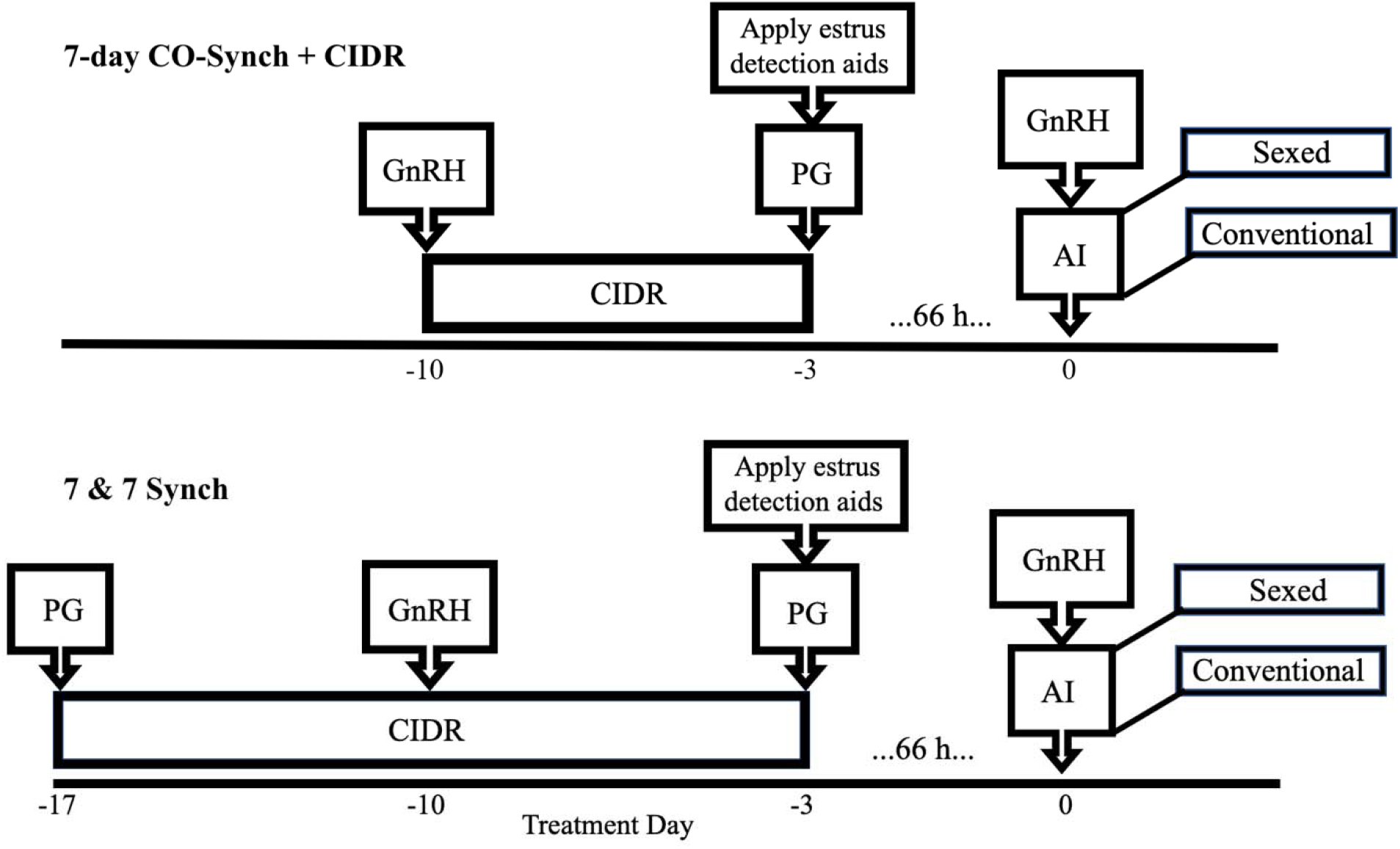
Treatment schedules for the 7-day CO-Synch + CIDR protocol with fixed-time AI (FTAI) and the 7 & 7 Synch protocol with FTAI. Cows treated with the 7-day CO-Synch + CIDR protocol (n = 769) received administration of 100 μg gonadotropin-releasing hormone (GnRH) and insertion of a 1.38 g intravaginal progesterone releasing insert (CIDR) on Day -10, and administration of 500 μg prostaglandin F2a (PG) coincident with CIDR removal on Day -3. Cows treated with the 7 & 7 Synch protocol (n = 769) received administration of PG and insertion of CIDR on Day -17, administration of GnRH on Day -10, and administration of PG coincident with CIDR removal on Day -3. Estrotect™ estrus detection aids were applied to all cows on Day -3 and patch activation score were recorded at the time of FTAI on Day 0.

### 2.2. Estrus Detection

Estrotect estrus detection aids (Estrotect™, Rockway Inc, Spring Valley, WI) were applied to all cows on Day -3 at the time of PG administration and CIDR removal, and patch activation was recorded at the time of FTAI on Day 0. Patch activation was scored on a numerical scale of 0 to 4 (0 = missing patch; 1 = 0-25% activated; 2 = 25-50% activated; 3 = 50-75% activated; and 4 = 75-100% activated) [13]. Estrus was defined as 50% or greater of the Estrotect patch activated or a missing patch (patch score 0, 3, 4).

### 2.3. Artificial Insemination

Within protocol in each location, cows were blocked based on age, DPP, and BCS and pre-assigned to receive either conventional or sex-sorted semen at FTAI (Figure 1). To minimize potential confounding of treatment effects in locations where FTAI was performed by two technicians, cows were also blocked on these criteria within protocol and semen type and pre-assigned to technician. Animals in Locations 1, 2, 4, 6, and 10 were inseminated by a single technician. Semen was collected from eight commercially available AI bulls, and units of sex-sorted and conventional semen were produced from contemporaneous ejaculates. Both conventional and sex-sorted semen from each bull passed the standard quality control criteria used for the respective semen types. Units of conventional semen were generated with 20.0 × 10^6^ live cells per 0.25 mL straw prior to freezing. Units of sex-sorted semen were produced using the SexedULTRA™ Genesis III sorting technology (Sexing Technologies, Navasota, TX) with 4.0 × 10^6^ live cells per 0.25 mL straw prior to freezing with a marketed level of > 90% accuracy for the desired sex. Sex-sorted units were sorted to either contain X (Bulls A, B, C, D, F, and G) or Y-bearing chromosomes (Bulls E and H). Bull numbers were minimized within location to limit variance associated with bull-to-bull fertility differences, with semen from one bull used within each location for Locations 2-11 and semen from two bulls used in Location 1. Semen from a total of eight bulls was used across the locations, as Locations 2, 3, 5, 10 and 11 used semen from Bull C.

### 2.4. Pregnancy Diagnosis

Pregnancy per AI (P/AI) was determined 75 to 90 days after artificial insemination by transrectal ultrasonography, using an Aloka 500V equipped with a 5.0-MHz linear-array transducer (Aloka, Wallingford, CT) in Locations 1-6 and a SonoSite EDGE equipped with a L52 10.0-5.0 MHz linear-array transducer (SonoSite, Inc., Bothwell, WA) in Locations 7-11.

### 2.5. Statistical Analysis

Statistical analyses were performed using SAS (SAS 9.4 Inst. Inc., Cary, NC). A general linear model (PROC GLM) was used to confirm balance across treatments with respect to DPP, BCS, and age of cows. Mixed models (PROC GLIMMIX) using the binomial distribution link logit function were used to evaluate the proportion of cows expressing estrus prior to FTAI and P/AI. Variables tested for inclusion in the mixed model for estrus expression were protocol (7 & 7 Synch or the 7-day CO-Synch + CIDR), age, BCS, DPP, and all two-way interactions. Location was included as a random effect. Estrous response by days postpartum and protocol was modeled from the regression equation coefficient estimates using the average body condition score (5.7) and age (4.6). Variables tested for inclusion in the mixed model for P/AI were protocol, semen type, age, BCS, DPP, and all two-way interactions. Bull, technician, location, and location × protocol × semen were included as random effects. Interactions not observed to have effect (*P* > 0.10) were removed from the models using backwards elimination, with the exception of the protocol × semen interaction central to the experiment. The final mixed model for P/AI included fixed effects of protocol, semen, age, BCS, DPP, and protocol × semen as well as the random effects of bull, technician, location, and location × protocol × semen. The three-way interaction of location × protocol × semen was included in the random statement to provide a more conservative analysis of P/AI, as treatment × location interaction is the most appropriate error term for tests of potential treatment differences in multi-location experiments. Models for P/AI were tested across all cows and were also tested specifically among cows that expressed estrus prior to FTAI as well as specifically among cows that failed to express estrus. Across treatments and within each treatment, Chi-square was used for comparisons of P/AI obtained among cows that expressed estrus vs. cows that failed to express estrus.

## 3. Results

### 3.1. Location Summary

Age, DPP, body condition score (BCS), and the proportion of cows expressing estrus prior to FTAI by treatment are summarized in Table 1. Age, DPP, and BCS did not differ between treatments within each location or across all locations. Individualized cow age was not known in Location 10, which consisted exclusively of mature cows seven years of age and older.

**Table 1.** Cow age, days postpartum (DPP), body condition score (BCS), and proportion expressing estrus prior to fixed-time artificial insemination (FTAI) by treatment and location.

| Treatment<sup>1</sup> | N | Age | DPP<sup>2</sup> | BCS<sup>3</sup> | Estrus expression prior to FTAI<sup>4</sup> |  |
| --- | --- | --- | --- | --- | --- | --- |
|  |  |  |  |  | Proportion | % |
| 7-day CO-Synch + CIDR | 769 | 4.6 ± 1.8 | 80.2 ± 19.3 | 5.7 ± 0.5 | 492/769 | 64<sup>b</sup> |
| Location 1 | 43 | 3.5 ± 1.7 | 98.8 ± 18.8 | 5.5 ± 0.5 | 31/43 | 72 |
| Location 2 | 72 | 4.1 ± 1.6 | 75.0 ± 11.8 | 5.7 ± 0.4 | 52/72 | 72 |
| Location 3 | 78 | 4.1 ± 0.4 | 98.3 ± 26.7 | 5.8 ± 0.1 | 59/78 | 76 |
| Location 4 | 59 | 4.8 ± 1.7 | 63.8 ± 17.1 | 5.8 ± 0.5 | 47/59 | 80 |
| Location 5 | 82 | 3.0 ± 1.6 | 76.2 ± 17.2 | 5.5 ± 0.4 | 56/82 | 68 |
| Location 6 | 94 | 5.3 ± 2.0 | 74.2 ± 11.8 | 5.9 ± 0.5 | 69/94 | 73 |
| Location 7 | 84 | 4.3 ± 1.2 | 66.6 ± 11.5 | 5.5 ± 0.4 | 35/84 | 42 |
| Location 8 | 58 | 5.1 ± 2.7 | 88.4 ± 16.1 | 5.9 ± 0.6 | 39/58 | 67 |
| Location 9 | 78 | 5.0 ± 0 | 82.0 ± 12.6 | 5.8 ± 0.6 | 41/78 | 53 |
| Location 10 | 62 | - | 87.1 ± 12.9 | 5.8 ± 0.4 | 32/62 | 52 |
| Location 11 | 59 | 6.0 ± 0 | 78.0 ± 13.9 | 5.7 ± 0.4 | 31/59 | 53 |
| 7 & 7 Synch | 769 | 4.6 ± 1.9 | 80.2 ± 19.1 | 5.8 ± 0.6 | 630/769 | 82<sup>a</sup> |
| Location 1 | 46 | 3.9 ± 1.8 | 95.2 ± 20.0 | 5.6 ± 0.6 | 41/46 | 89 |
| Location 2 | 71 | 4.1 ± 1.4 | 73.4 ± 12.9 | 5.6 ± 0.4 | 56/71 | 79 |
| Location 3 | 91 | 4.8 ± 2.1 | 94.0 ± 26.1 | 5.8 ± 0.4 | 85/91 | 93 |
| Location 4 | 58 | 4.5 ± 1.6 | 64.2 ± 16.8 | 6.1 ± 0.7 | 49/58 | 85 |
| Location 5 | 77 | 3.0 ± 1.6 | 76.3 ± 17.2 | 5.5 ± 0.4 | 56/77 | 73 |
| Location 6 | 96 | 5.3 ± 1.9 | 74.2 ± 11.8 | 5.9 ± 0.5 | 84/96 | 88 |
| Location 7 | 71 | 4.3 ± 1.2 | 66.6 ± 11.5 | 5.5 ± 0.4 | 47/71 | 66 |
| Location 8 | 61 | 5.1 ± 2.7 | 88.4 ± 16.1 | 5.9 ± 0.6 | 55/61 | 90 |
| Location 9 | 74 | 5.0 ± 0 | 82.0 ± 12.7 | 5.8 ± 0.6 | 54/74 | 73 |
| Location 10 | 63 | - | 87.1 ± 12.9 | 5.7 ± 0.4 | 52/63 | 83 |
| Location 11 | 61 | 6.0 ± 0 | 78.8 ± 14.0 | 5.7 ± 0.4 | 51/61 | 84 |

Data presented as mean value (± standard deviation of mean)
<sup>1</sup>See Figure 1 for treatment descriptions.
<sup>2</sup>DPP calculated from calving date to FTAI date
<sup>3</sup>BCS of cows recorded on Day -17 using a scale of 1-9 (1 = emaciated and 9 = obese)
<sup>4</sup>Proportion of cows expressing estrus prior to FTAI, based on ≥ 50% of the patch coating rubbed off or a missing patch
<sup>ab</sup>Values with different superscripts differ ( $P = 0.01$ )

### 3.2. Estrus Expression

The proportion of cows expressing estrus prior to FTAI is summarized in Table 1. The proportion of cows expressing estrus was increased (*P* = 0.01) among cows treated with 7 & 7 Synch (82%; 630/769) compared with the 7-day CO-Synch + CIDR (64%; 492/769). Estrus expression was also affected by a protocol × DPP interaction (*P* = 0.0004), with 7 & 7 Synch resulting in a greater magnitude increase in the proportion of cows expressing estrus prior to FTAI among cows with greater DPP. To visualize the protocol × DPP interaction, regression equation coefficient estimates from the GLIMMIX procedure of SAS (SAS 9.4 Inst. Inc., Cary, NC) were used to model the effect of protocol across the DPP range. Figure 2 represents modeled likelihood of estrus expression assuming the average body condition score (5.7) and age (4.6) across cows in the experiment. The proportion of cows expressing estrus prior to FTAI was also affected by BCS (*P* = 0.05), with an increased proportion of cows in greater BCS expressing estrus. Additionally, estrus expression tended to be affected by a protocol × age interaction (*P* = 0.08), with 7 & 7 Synch resulting in a greater magnitude increase among older cows with respect to the proportion of cows expressing estrus prior to FTAI.

**Figure 2.**
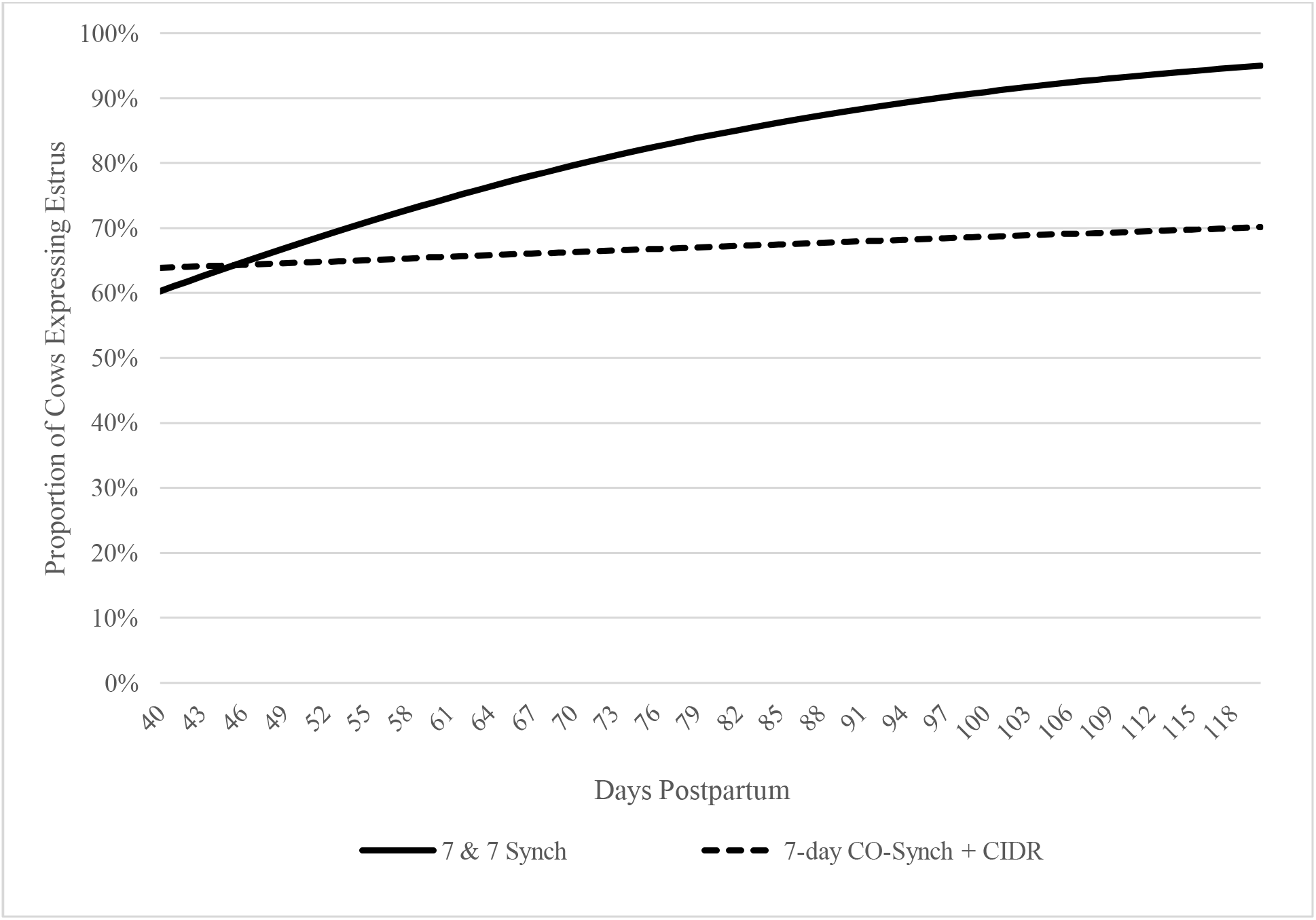
Modeled proportion of cows expressing estrus prior to fixed-time artificial insemination (FTAI) based on days postpartum (DPP) and protocol. Estrus expression was affected by a protocol × DPP interaction (*P* = 0.0004), with 7 & 7 Synch resulting in a greater increase in the proportion of cows expressing estrus prior to FTAI among cows with greater DPP. A mixed model was generated using regression equation coefficient estimates from the GLIMMIX procedure of SAS (SAS 9.4 Inst. Inc., Cary, NC). The model contained effects of protocol, DPP, body condition score, age and DPP × protocol. Figure represents modeled likelihood of estrus expression assuming the average body condition score (5.7) and age (4.6) across cows in the experiment.

### 3.3. Pregnancy Rate

Pregnancy outcomes are summarized by location in Table 2. Pregnancy rates were affected by protocol (*P* = 0.001), semen type (*P* < 0.0001), age (*P* = 0.04), and DPP (*P* = 0.02). Regardless of semen type used for AI, P/AI was increased (*P* = 0.001) among cows treated with 7 & 7 Synch (conventional semen: 72% [280/389]; sex-sorted semen: 52% [199/380]) compared with 7-day CO-Synch + CIDR (conventional semen: 60% [231/383]; sex-sorted semen: 44% [171/386]). Irrespective of treatment, sex-sorted semen resulted in decreased (*P* < 0.0001) P/AI compared with conventional semen. Pregnancy results based on protocol, semen type, and expression of estrus prior to FTAI are presented in Table 3. Across treatments and within each treatment, cows that expressed estrus prior to FTAI achieved greater P/AI (*P* < 0.001). In comparison with conventional semen, sex-sorted semen resulted in reduced P/AI both among cows that expressed estrus (*P* < 0.0001) and among cows that failed to express estrus (*P* = 0.001) prior to FTAI. Specifically among cows that expressed estrus prior to FTAI, greater P/AI (*P* = 0.01) was achieved among cows receiving 7 & 7 Synch compared with cows receiving the 7-day CO-Synch + CIDR protocol. Among cows that failed to express estrus prior to FTAI, pregnancy rates did not differ significantly (*P* = 0.14) between protocols at this power of test.

**Table 2.** Pregnancy rates (P/AI)^1^ to fixed-time artificial insemination^2^

| Treatment<sup>3</sup> | P/AI<br>Conventional Semen |  | P/AI<br>Sex-Sorted Semen |  |
| --- | --- | --- | --- | --- |
|  | Proportion | % | Proportion | % |
| 7-day CO-Synch + CIDR | 233/383 | 61<sup>b</sup> | 171/386 | 44<sup>d</sup> |
| Location 1 | 15/21 | 71 | 15/22 | 68 |
| Location 2 | 24/37 | 65 | 13/35 | 37 |
| Location 3 | 24/40 | 60 | 17/38 | 45 |
| Location 4 | 18/30 | 60 | 14/29 | 48 |
| Location 5 | 20/40 | 50 | 14/42 | 33 |
| Location 6 | 29/47 | 62 | 18/47 | 38 |
| Location 7 | 24/40 | 60 | 20/44 | 46 |
| Location 8 | 19/30 | 63 | 15/28 | 54 |
| Location 9 | 22/37 | 59 | 22/41 | 54 |
| Location 10 | 19/31 | 61 | 13/31 | 42 |
| Location 11 | 19/30 | 63 | 10/29 | 35 |
| 7 & 7 Synch | 280/389 | 72<sup>a</sup> | 199/380 | 52<sup>c</sup> |
| Location 1 | 19/25 | 76 | 13/21 | 62 |
| Location 2 | 26/36 | 72 | 20/35 | 57 |
| Location 3 | 38/45 | 84 | 25/46 | 54 |
| Location 4 | 22/29 | 76 | 11/29 | 38 |
| Location 5 | 27/39 | 69 | 17/38 | 45 |
| Location 6 | 30/48 | 63 | 28/48 | 58 |
| Location 7 | 22/36 | 61 | 16/35 | 46 |
| Location 8 | 23/31 | 74 | 20/30 | 67 |
| Location 9 | 26/37 | 70 | 21/37 | 57 |
| Location 10 | 25/31 | 81 | 12/32 | 38 |
| Location 11 | 22/32 | 69 | 16/29 | 55 |

<sup>1</sup>P/AI was determined via transrectal ultrasonography 80-85 days after AI<sup>2</sup>Fixed-time artificial insemination was performed 66 hours after administration of PG<sup>3</sup>See Figure 1 for treatment descriptions<sup>abcd</sup>Values with different superscripts differ ( $P = 0.001$ )

**Table 3.** Pregnancy rates to fixed-time artificial insemination (FTAI) based on treatment, semen type, and expression of estrus prior to FTAI

| Treatment<sup>1</sup> | Conventional Semen |  |  |  | Sex-Sorted Semen |  |  |  | Across Semen Types |  |  |  |
| --- | --- | --- | --- | --- | --- | --- | --- | --- | --- | --- | --- | --- |
|  | P/AI |  | P/AI |  | P/AI |  | P/AI |  | P/AI |  | P/AI |  |
|  | Estrous<sup>4</sup> Cows |  | Non-estrous<sup>5</sup> Cows |  | Estrous Cows |  | Non-estrous Cows |  | Estrous Cows |  | Non-Estrous Cows |  |
|  | Proportion | % | Proportion | % | Proportion | % | Proportion | % | Proportion | % | Proportion | % |
| 7-day CO-Synch + CIDR | 170/255 | 67 | 63/128 | 49 | 120/237 | 51 | 50/149 | 34 | 290/492 | 59<sup>b</sup> | 113/277 | 41<sup>c</sup> |
| 7 & 7 Synch | 243/322 | 75 | 37/67 | 55 | 171/308 | 55 | 28/72 | 38 | 414/630 | 66<sup>a</sup> | 65/139 | 47<sup>c</sup> |

<sup>1</sup>See Figure 1 for treatment descriptions
<sup>2</sup>Pregnancy rate to AI was determined via transrectal ultrasonography 80-85 days after FTAI
<sup>3</sup>FTAI was performed 66 hours after administration of PG (Day -17) by a pre-assigned technician
<sup>4</sup>Estrous cows based on $\geq 50\%$ of the patch coating rubbed off or a missing patch at FTAI
<sup>5</sup>Non-estrous was recorded as having $< 50\%$ of the patch coating rubbed off at FTAI
<sup>abcd</sup>Values with different superscripts differ ( $P = 0.01$ )

## 4. Discussion

7 & 7 Synch improved pregnancy outcomes among postpartum beef cows receiving FTAI in comparison to results obtained using the 7-day CO-Synch + CIDR protocol. By significantly increasing the proportion of cows that express estrus prior to FTAI, a significant increase in P/AI was obtained using 7 & 7 Synch. Furthermore, P/AI was greater following 7 & 7 Synch whether using conventional or sex-sorted semen. The significant effect of protocol on the proportion of cows expressing estrus as well as on P/AI was observed across cows of varying age, DPP, and BCS. However, the magnitude of improvement observed in the proportion of cows expressing estrus was greater among cows in greater DPP ranges and tended to be greater among cows of increased age. Given the relationship of age and DPP with estrous cyclicity, this observation supports the hypothesized mechanism for improved control of the estrous cycle using 7 & 7 Synch. Induced persistence of the dominant follicle, achieved via treatment with PG and a CIDR insert in advance of GnRH, should only occur among cows that have already initiated estrous cyclicity after calving.

Short-term estrus synchronization protocols such as the 7-day CO-synch + CIDR rely on administration of exogenous GnRH at the beginning of the protocol to induce formation of a new follicular wave. However, as a consequence of animals beginning an estrus synchronization protocol at varying stages of the estrous cycle, a proportion of animals are in a stage of follicular development that do not have the capacity to respond to the initial GnRH administration. As a result, a proportion of females undergo luteolysis during the course of CIDR treatment, thereby developing a persistent, subfertile follicle that is maintained until CIDR removal [14,15]. Although these follicles will appear physiologically mature, the oocyte is functionally compromised due to intrafollicular changes resulting in premature resumption of meiosis [16]. The single-step approach of combining a progestin and PG at the beginning of 7 & 7 Synch induces persistent follicle formation among a large proportion of cows in advance of GnRH administration [8]. However, the intentional formation of persistent follicles is advantageous in this context as a means to enhance follicular maturity on Day -10 [8], thereby improving likelihood of ovulatory response to GnRH administration and subsequent emergence of a new synchronized follicular wave.

In the present experiment, the increase observed in the proportion of cows that expressed estrus prior to FTAI following 7 & 7 Synch is attributed to the greater degree of uniformity previously observed among cows using this treatment schedule [8]. By improving follicular maturity in advance of GnRH administration, variation among cows is reduced later in the protocol. At the time of PG administration on Day -3, a greater proportion of cows were observed to have a single CL following treatment with 7 & 7 Synch, whereas a greater proportion of cows were observed to have no CL or a CL and an accessory CL following treatment with the 7-day CO-Synch + CIDR [8]. Recognizing that 7 & 7 Synch induces luteolysis among the majority of cyclic cows via PG administration on Day 0, the observation that the majority of cows presented with a single CL on Day -3 indicates a successful response to GnRH administration on Day -10 and an enhanced degree of uniformity among the synchronized group.

The proportion of cows expressing estrus prior to fixed-time AI was greatly increased among cows treated with the 7 & 7 Synch protocol. Results from this field trial emphasize the importance of estrus expression prior to FTAI, as the published literature consistently notes that P/AI is increased among cows who express estrus compared with cows who fail to express estrus prior to FTAI [17]. In the present study, 82% (630/769) of cows treated with the 7 & 7 Synch expressed estrus prior to FTAI while 64% (492/769) of the cows treated with the 7-day CO-Synch + CIDR expressed estrus prior to FTAI. Correspondingly, P/AI was increased among cows that expressed estrus prior to FTAI compared with cows that failed to express estrus at FTAI. Among cows treated with 7 & 7 Synch, cows that expressed estrus had an increased pregnancy rate (conventional semen: 75% [243/322]; sex-sorted semen: 55% [171/308]) compared with non-estrous cows (conventional semen: 55% [37/67]; sex-sorted semen: 38% [28/72]). Likewise, among cows treated with the 7-day CO-Synch + CIDR, cows that expressed estrus also obtained increased P/AI (conventional semen: 67% [170/255]; sex-sorted semen: 51% [120/237]) compared with non-estrous cows (conventional semen: 49% [63/128]; sex-sorted semen: 34% [50/149]). While estrus expression was associated with greater pregnancy rates in both protocols, a greater proportion of cows expressed estrus prior to FTAI following 7 & 7 Synch, resulting in greater overall P/AI.

Moreover, when the analysis was restricted only to cows that expressed estrus prior to FTAI, greater P/AI (*P* = 0.01) was achieved among estrous cows receiving 7 & 7 Synch compared with estrous cows receiving the 7-day CO-Synch + CIDR. This merits further investigation, as this observation suggests an amelioration of subfertility even among cows that express estrus prior to FTAI. In the standard 7-day CO-Synch + CIDR protocol, subfertility may result among cows that failed to ovulate in response to the initial administration of GnRH and underwent luteolysis prior to CIDR removal. This subpopulation of cows ovulates a persistent follicle, resulting in subfertility despite an otherwise normal display of standing estrus [14–16]. In theory, considering the treatment schedule of 7 & 7 Synch, potential for subfertility stemming from ovulation of a persistent follicle could be reduced or eliminated. An alternative or additional explanation for improved fertility observed specifically among estrous cows could stem from optimized timing of ovulation relative to FTAI following 7 & 7 Synch. In a previous large multi-location embryo transfer field trial, 7 & 7 Synch resulted in a greater proportion of cows expressing estrus earlier in the distribution of estrus onset [9]. This observation suggests FTAI would be occurring later relative to onset of estrus on average and therefore closer to the timing of ovulation. Timing of insemination clearly affects fertility, with some indication in the literature that insemination closer to ovulation is particularly critical for some bulls [18–21]. Further investigation may be needed to assess the degree to which these factors may contribute to the significant increase in P/AI observed with 7 & 7 Synch among cows that expressed estrus prior to FTAI.

The significant increase observed in the proportion of cows expressing estrus prior to FTAI following 7 & 7 Synch suggests a practical opportunity to reduce incidence of embryonic loss. Previous investigations have observed a reduction in pregnancy loss among cows that express estrus near the time of AI [13,22,23]. Animals that express estrus within 24 hours of FTAI have increased diameter dominant follicles and increased serum estradiol concentrations, corresponding to an increase in pregnancy success [24]. Estradiol produced by the dominant follicle is critical for regulation of physiological processes that aid in establishment and maintenance of pregnancy in cattle [25]. With this understanding, maximizing the number of animals that express estrus and present with a physiologically mature preovulatory follicle at the time of FTAI is critical [26]. The proportion of cows expressing estrus following 7 & 7 Synch and the follicular maturity at the time of FTAI in previous studies [8] raise questions as to whether reduced rates of embryonic loss are driving the enhanced fertility observed with this protocol.

Although pregnancy rates were increased among cows treated with 7 & 7 Synch across semen types, use of sex-sorted semen resulted in reduced pregnancy rates in comparison with conventional semen irrespective of protocol. Despite the numerous hypothetical benefits that progeny sex selection offers in the beef industry, adoption of sex sorted semen has been limited by the increased cost per unit of sex-sorted semen and the decreased success in FTAI programs semen [27,28]. Lower pregnancy rates have been observed across the literature when sex-sorted semen is used in FTAI programs [29–31]. The sex-sorting procedure contributes to damage, decreased quality and quantity of cells, as well as pre-capacitation and a reduced fertility lifespan of sex-sorted sperm in the female reproductive tract [30]. Some investigators have suggested that sex-sorted semen has increased sensitivity to the timing of insemination in relation to ovulation and onset of estrus [32]. Cows that fail to express estrus prior to FTAI have particularly decreased P/AI when sexed semen is used, perhaps due to the reduced sperm lifespan and timing of ovulation not being optimally aligned [29]. With this understanding, P/AI with sex-sorted semen is optimized in females when estrus is expressed prior to FTAI [27,33,34]. An estrus synchronization protocol that optimizes estrus expression among females prior to FTAI may increase utilization of sex-sorted semen across operations in the beef industry. In the present experiment, 7 & 7 Synch significantly increased both the proportion of cows expressing estrus and the P/AI with sex-sorted semen when compared with the 7-day CO-Synch + CIDR.

Another factor that can alter success of synchronization and P/AI in postpartum beef cows is the proportion of cows that are anestrous at the start of the breeding season [35]. Factors that influence the proportion of cows that resume estrous cyclicity prior the beginning of the breeding season include DPP, parity and BCS [36]. Across protocols, BCS had a significant effect (*P* = 0.05) on the proportion of cows expressing estrus, with a greater proportion of cows in greater body condition expressing estrus prior to FTAI. Cows that are in body condition scores less than 5 have an increased potential for an extended period of anestrus when compared with cows in greater BCS [36]. Body condition score greater than 5 at calving is positively correlated with increased follicular development and pituitary LH content at 30 days after calving, physiological mechanisms which are essential in resumption of estrous cyclicity [37].

Several observations from the present study suggest the enhanced P/AI observed following 7 & 7 Synch is primarily due to an effect specific to cows that are cycling at the start of the protocol. For example, there was a significant effect of DPP × protocol (Figure 2) on the proportion of cows expressing estrus prior to FTAI, with 7 & 7 Synch resulting in a greater magnitude of improvement among cows with greater DPP. There is a positive correlation between DPP and cyclicity; as DPP increases, cows are more likely to have resumed normal estrous cycles compared with cows with shorter DPP [35]. Likewise, the proportion of cows expressing estrus prior to FTAI tended to be affected by age × protocol (*P* = 0.08), with 7 & 7 synch resulting in a greater magnitude of improvement among cows of greater age. This tendency could also support the hypothesis that improvements resulting from 7 & 7 Synch stem primarily from an effect observed among cows that are cyclic at the initiation of the protocol. Nutrient use for maintenance requirements, basic energy reserves, activity, and growth are prioritized over reproduction [36]. Primiparous cows have increased nutrient demand due to their additional growth requirements, resulting in prioritization of nutrient use for growth rather than reproduction. The low physiological prioritization of reproductive processes increases the length of anestrus and limits the proportion of primiparous cows that have resumed normal estrous cyclicity at the start of the breeding season [36,38]. Thus, the interaction of protocol and age observed in this experiment supports the hypothesis that the treatment schedule of 7 & 7 Synch is primarily advantageous for cows that have resumed estrous cyclicity prior to initiation of treatment.

When compared with the 7-day CO-Synch + CIDR protocol, 7 & 7 Synch resulted in a greater proportion of cows expressing estrus prior to FTAI and enhanced P/AI to both conventional and sexed semen. Resultantly, 7 & 7 Synch provides a promising opportunity to increase utilization of sexed semen in the beef industry as well as improve success rates with FTAI regardless of semen type. The increased proportion of cows expressing estrus prior to FTAI are associated with greater P/AI, however the significant increase in P/AI among cows treated with 7 & 7 Synch merits further mechanistic investigation. With only one more additional handling of animals and one additional week in length of protocol compared with the 7-day CO-Synch + CIDR, 7 & 7 Synch offers much potential as a platform to improve success with fixed-time AI among postpartum beef cows.

## 5. Acknowledgments

We gratefully acknowledge and are deeply appreciative of the following companies and farms who allowed this project to be possible: Sexing Technologies (Navasota, Texas) for providing semen and funding; Merck Animal Health (Madison, New Jersey) for providing Fertagyl and Estrumate; Zoetis (Madison, New Jersey) for providing EAZI-Breed CIDR cattle inserts; Estrotect Inc. (Spring Valley, Wisconsin) for providing estrus detection aids; and the Mason-Knox Ranch (Frankfort, South Dakota), D&R Ogren Farm (Langford South Dakota), Adrian Farms (Milan, Missouri), Willard Farms (Sleeper, Missouri) and the University of Missouri Southwest Research Center (Mount Vernon, Missouri) for the use of animals in this project.

